# Beet taproot plasma membrane sugar transport revisited

**DOI:** 10.1101/2021.09.21.461191

**Authors:** Antonella Reyer, Nadia Bazihizina, Justyna Jaślan, Sönke Scherzer, Nadine Schäfer, Dawid Jaślan, Dirk Becker, Thomas D. Müller, Benjamin Pommerrenig, H. Ekkehard Neuhaus, Irene Marten, Rainer Hedrich

## Abstract

Sugar beet (*Beta vulgaris*) is the major sugar-producing crop in Europe and Northern America, as the taproot stores sucrose at a concentration of around 20%. Genome sequence analysis together with biochemical and electrophysiological approaches led to the identification and characterization of the TST sucrose transporter driving vacuolar sugar accumulation in the taproot. However, the sugar transporters mediating sucrose uptake across the plasma membrane of taproot parenchyma cells remained unknown. As with glucose, sucrose stimulation of taproot parenchyma cells caused inward proton fluxes and plasma membrane depolarization, indicating a sugar/proton symport mechanism. To decipher the nature of the corresponding proton-driven sugar transporters, we performed transcriptomic taproot profiling and identified the cold-induced PMT5a and STP13 transporters. When expressed in *Xenopus laevis* oocytes, BvPMT5a was characterized as a voltage- and H^+^-driven low-affinity glucose transporter, which does not transport sucrose. In contrast, BvSTP13 operated as a high-affinity H^+^/sugar symporter, transporting glucose better than sucrose, and being more cold-tolerant than BvPMT5a. Modeling of the BvSTP13 structure with bound mono- and disaccharides suggests plasticity of the binding cleft to accommodate the different saccharides. The identification of BvPMT5a and BvSTP13 as taproot sugar transporters could improve breeding of sugar beet to provide a sustainable energy crop.

**Significance Statement:** *In vivo* electrophysiological studies with sugar beet taproots provide clear evidence for proton-coupled glucose and sucrose uptake into taproot parenchyma cells. In search for the molecular entities, the taproot-expressed BvPMT5a and BvSTP13 carriers were studied in detail, because they mediate proton-driven import of glucose and sucrose and thus provide proper candidates for sugar beet plasma membrane sugar-proton cotransporters.

## INTRODUCTION

Sugar beet (*Beta vulgaris*) and sugarcane (*Saccharum officinarum*) together account for the world’s sugar production. Industrial production of sugar from the sugar beet taproot began in the early nineteenth century and since then, breeding has increased the sugar content from 8% to about 21%. From the late 1970s to the present, plant scientists have been trying to identify the transport proteins that translocate sucrose from the sugar factories in the leaf to its final depot in the taproot. In the early days of sugar beet (*Beta vulgaris*) research, sucrose transport was examined in taproots, leaf discs and plasma membrane-enriched vesicles (Wyse, 1979; Bush, 1989; Sakr et al., 1993). Translocation of sucrose against its concentration gradient requires metabolic energy. The first *in vitro* evidence for a proton-driven sucrose symport, which uses the proton motive force (PMF) as a secondary energy source for solute uphill transport, was provided by Bush from studies on plasma membrane vesicles from sugar beet leaves (1989). Sucrose import was also found very sensitive to changes in electrical membrane potential as well as to orthovanadate, an inhibitor of the H^+^-ATPase, whose H^+^ pumping ability is required to keep the resting membrane voltage hyperpolarized, and to retain the inward H^+^ gradient (Buckhout, 1989; Bush, 1990; Michonneau et al., 2004).

With the beginning of the molecular era, the nature of the first plant glucose and sucrose transporters was identified (Sauer & Tanner, 1989; Sauer et al., 1990; Riesmeier et al., 1992). In 2013, the genome of *Beta vulgaris* was sequenced (Dohm et al., 2014), providing access to the molecular transporter inventory of sugar beet. Among them, BvSUT1 was demonstrated to represent a proton-driven transporter sucrose loader of the sugar beet phloem in source leaves (Nieberl et al., 2017), while the BvTST2.1 transporter was shown to be responsible for vacuolar sucrose accumulation in the sugar beet taproots (Jung et al., 2015). In patch-clamp studies with vacuoles of BvTST2.1-overexpressing tobacco mesophyll cells, this sucrose-specific transporter was characterized as a proton/sucrose antiporter, which couples the import of sucrose into the vacuole with the export of protons (Jung et al., 2015). Release of vacuolar sucrose and glucose is mediated by sugar symporters of the BvSUT4/AtSUC4 and BvIMP/AtERDL6 types (Schulz et al., 2011; Schneider et al., 2012; Klemens et al., 2014; Rodrigues et al., 2020).

Plants respond to cold temperatures by accumulating compatible solutes (e.g., proline, polyols, soluble sugars such as sucrose, glucose, fructose, raffinose) to protect cellular integrity (Wanner & Junttila, 1999; Gusta et al., 2004; Krasensky & Jonak, 2012; Klemens et al., 2014; Tarkowski & Van den Ende, 2015; Keller et al., 2021). Interestingly, altered vacuolar glucose levels are accompanied by changes in vacuolar sugar transporter gene activities. In Arabidopsis plants, transcript levels of the vacuolar glucose unloader AtERDL6 and vacuolar glucose loader AtTST1/2 were shown sensitive to cold exposure (Poschet et al., 2011). The latter observation is in agreement to results from studies with AtTST1 loss-of-function and BvIMP-overexpressing Arabidopsis plants, suggesting that the ability of the plant to accumulate monosaccharides in vacuoles under cold conditions accounts for frost tolerance (Klemens et al., 2014). In sugar beet taproots, the transcription of the vacuolar sucrose loader BvTST2.1 and sucrose unloader BvSUT4 changed when exposed to low temperatures (Rodrigues et al., 2020), a finding that may be related to the cold sensitivity of the sugar beet plant.

In this sugar beet study, we were especially interested in plasma membrane sugar transporters that could contribute to apoplastic loading and accumulation of sugars in the taproot. Using voltage-recording and proton-sensing microelectrodes, we directly monitored sucrose- and glucose-dependent changes in the membrane potential and proton flux in taproot slices. Together these findings provide the first *in vivo* evidence for proton-coupled sugar uptake in taproot parenchyma cells. In search of involved transporters, we identified the cold-regulated BvPMT5a and BvSTP13 transporters. Based on their transport features, we shortlisted both BvPMT5a and BvSTP13 as potential candidates for taproot plasma membrane proton-coupled glucose and sucrose uptake.

## RESULTS

### Sugar uptake in taproots is directly linked to proton influx and membrane depolarization

As soon as sucrose is translocated from the source leaves to the taproot and released from the phloem to the apoplast, sucrose needs to enter the storage parenchyma cells (Lemoine et al., 1988; Godt & Roitsch, 2006). For this, sucrose is most likely translocated across the plasma membrane via H^+^-coupled sugar uploaders. To date, insights into *Beta vulgaris* taproot sugar transport have mostly been gained by classical physiological assays such as uptake of radioactive sugars into tissue slices (*in vivo*) and plasma membrane enriched vesicles (*in vitro*). Based on the ground-breaking findings on *Beta vulgaris* sugar transport gained with taproot slices, we took advantage of the same experimental system. To first visualize cellular sucrose loading in the storage organ, we employed the fluorescent sucrose analog esculin, which is a substrate of several sucrose transporters (Gora et al., 2012; Reinders et al., 2012; Abelenda et al., 2019). After incubation of taproot slices from maturating 14 to 18-week-old sugar beets, strong fluorescent signals associated with the nucleus of parenchyma cells were detected (Figure S1). This finding is in agreement with esculin being a substrate of plasma membrane sugar transporters.

Theoretically, sugars can enter target cells via secondary active and/or passive transport. To explore the energetics of sucrose and glucose uptake into the taproot parenchyma, we investigated the corresponding plasma membrane electro-chemistry using voltage-recording and ion-selective electrodes. In the case of uphill plasma membrane sugar transport, the influx of the sugar is coupled to the down-hill flux of the proton (see for review Geiger 2011). To first monitor sucrose- and glucose-induced changes in H^+^ fluxes across the plasma membrane of taproot cells non-invasively and online, we employed scanning H^+^-selective electrodes (cf. Reyer et al., 2020). The parenchyma cells in taproot slices of 14- to 16-week-old sugar beets operated a H^+^ efflux before sugar stimulation (Fig 1a). This resting H^+^ efflux most likely reflects the H^+^ pump activity of the plasma membrane P-type ATPase, required to keep the membrane potential in its hyperpolarized state (Figure 1b). In line with a proton-coupled sugar symporter, administration of both glucose and sucrose (50 mM) resulted in a pronounced, transient decrease in the H^+^ efflux often lasting at least 30 minutes (Figure 1a, Figure S2a). During this glucose and sucrose provoked phase, maximum proton fluxes of 25.8 ± 7.2 nmol m^-2^ s^-1^ (n = 11, SEM) and 24.1 ± 7.4 nmol m^-2^ s^-1^ (n = 11, SEM) respectively, were measured. This response indicated that the plasma membrane of the parenchyma cells from slices derived from sugar-accumulating taproots is competent for sucrose and glucose transport.

**Figure 1.**
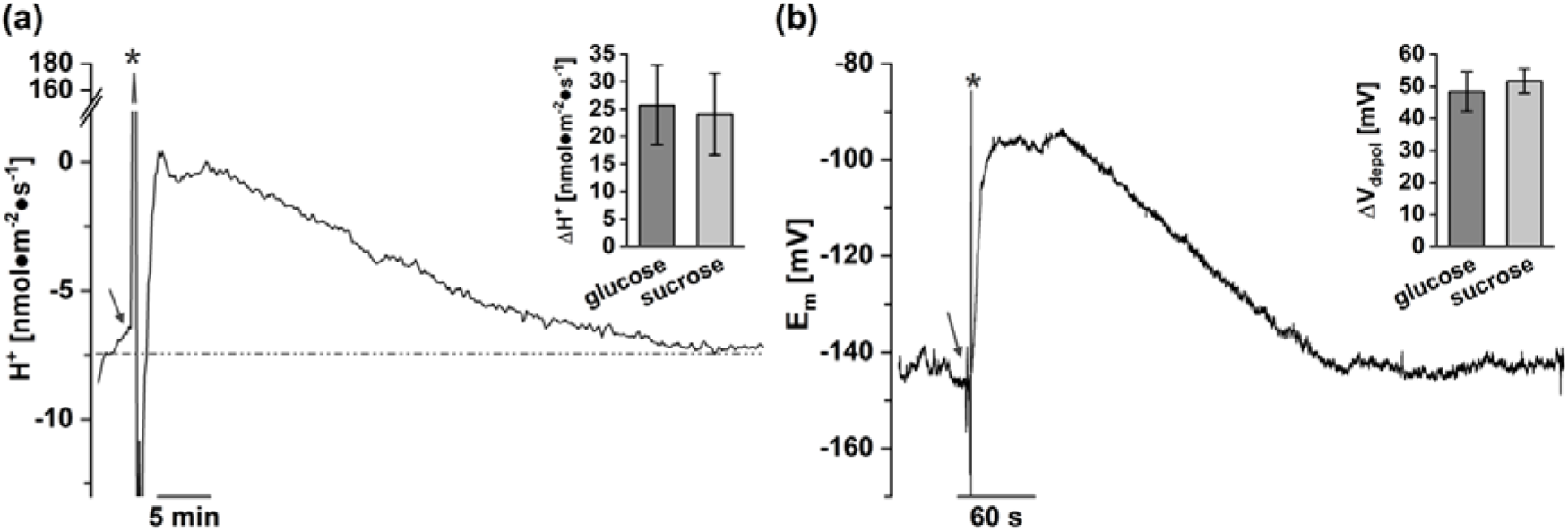
Glucose- and sucrose-induced changes in H^+^ fluxes and membrane depolarization of *Beta vulgaris* taproot cells. (a) H^+^ flux trace recorded from taproot slices in response to 50 mM glucose treatment. Time of glucose application is denoted by the arrow. Negative and positive fluxes represent H^+^ release from the cells and H^+^ uptake into the cells, respectively. The H^+^ flux level determined at rest shortly before sugar administration is indicated by a dotted line. The bar graph shows the maximal glucose- and sucrose-induced changes in the H^+^ fluxes relative to the H^+^ flux level at rest. Data represent means ± SEM with n = 11 each for glucose and sucrose. (b) Free running membrane voltage trace recorded from a taproot slice in response to 50 mM glucose. Time of glucose application is indicated by the arrow. The bar graph shows the maximal glucose- and sucrose-induced depolarization of the slices. Data represent means ± SEM of seven or six different taproots for glucose and sucrose, respectively. In a, b, taproots from GT2 were used. Grey stars (*) mark artefacts due to electrical disturbance during glucose application.

In plant cells, the plasma-membrane proton efflux results from the H^+^ pump activity of vanadate-sensitive AHA-type H^+^-ATPases (cf. Reyer et al., 2020, and references therein). Since sugars are not charged while protons are, one must predict that the phenomenon observed for the H^+^ fluxes (Figures 1a, S2a) is best explained mechanistically by H^+^/sugar co-import. To monitor sugar-induced membrane potential changes, parenchyma cells of the afore identified sugar-sensitive taproot slices were impaled with voltage recording microelectrodes. The membrane potential was −149.9 ± 3.3 mV (n = 18, SEM) at rest, and transiently depolarized by 48.3 ± 6.1 mV (n = 7, SEM) and 51.6 ± 3.7 mV (n = 6, SEM) upon addition of 50 mM glucose and sucrose, respectively (Figures 1b, S2b). Without removal of sugar, the membrane voltage generally slowly relaxed to the pre-stimulus level; a behavior in line with a depolarization and H^+^ influx-dependent activation of the H^+^-ATPase (Reyer et al., 2020). To exclude the possibility of an osmotic rather than sugar transport specific effect, we applied mannitol at same concentration. Mannitol, however, did not cause a membrane depolarization (Fig. S3).

### Search for taproot genes encoding glucose and sucrose transporters

Our electrophysiological studies with taproot slices point to a proton-pumping moiety in sugar-accumulating taproot cells that drives H^+^-coupled sucrose and glucose transport. These findings raise questions about the molecular nature and sugar specificity of the transporters involved. Members of the SUT family such as BvSUT3 and BvSUT1 likely play a pivotal role in sucrose uptake in sugar beet taproots (Fig. S5a). However, SUT transporters accept only sucrose and exclude monosaccharides for transport (Reinders et al., 2012). Given that we observed both plasma membrane depolarization and proton influx upon sucrose and glucose stimulation of taproot parenchyma cells (Figures 1, S2), it is reasonable to assume that other transporter, most likely members from the <u>M</u>ono<u>s</u>accharide <u>T</u>ransporter (MST) superfamily (Büttner, 2007; Pommerrenig et al., 2018), could also provide likely candidates for proton-coupled sugar transport. Of the seven MST subfamilies, only the INT, STP and PMT subfamilies harbor plasma membrane transporters (Scholz-Starke et al., 2003; Klepek et al., 2005; Schneider et al., 2006; Schneider et al., 2007; Klepek et al., 2010; Rottmann et al., 2018). INT subfamily members have been reported as carriers for inositol and other polyols, but not for sucrose (Schneider et al., 2006; Schneider et al., 2007). In contrast, STPs and PMTs exhibit a high specificity for the monosaccharides glucose and fructose (Klepek et al., 2005; Rottman et al., 2018). Astonishingly, the MdSTP13a homolog from apple (*Malus domesticus*), in contrast to the Arabidopsis homologue AtSTP13, was recently shown to transport the disaccharide sucrose and its analog esculin in addition to the monosaccharide glucose (Li et al., 2020). Therefore, we focused our search on the STP and PMT subfamilies. They consist of 14 putative STP and five PMT transport proteins (Figure S4). Among the expressed STP transporters (Figure S5b) we selected BvSTP13 because it is a homologue of apple MdSTP13 (Li et al. 2020), which was associated with the transport of several monosaccharides and sucrose. Among the PMTs, BvPMT5a and BvPMT5b transporters were the most highly expressed plasma membrane PMT transporter (Fig. S5b). We focused now on BvPMT5a because its transcript level, like that of BvSTP13, increased markedly in taproots when exposed to low temperatures, an important abiotic stress factor in sugar beet cultivation (Fig. S5b, c).

### BvPMT5a is a proton-coupled glucose and polyol transporter

To gain insights into the possible role of BtPMT5a in taproot sugar uptake and in relation to the *in vivo* electrophysiological recordings from the taproot slices (Figures 1, S2), the molecular transport characteristics of BvPMT5a were analyzed in *Xenopus laevis* oocytes via voltage-clamp recordings. This heterologous expression system is well accepted in the field of animal and plant ion transporter and also enables the measurement of H^+^-coupled transport of uncharged molecules such as sugars (Carpaneto et al., 2005; Nieberl et al., 2017). When BvPMT5a-expressing oocytes were exposed to the disaccharide sucrose (10 mM), no additional current to the current background noise was elicited (Figure 2b). However, application of the monosaccharides glucose or fructose arising from invertase-mediated sucrose breakdown caused pronounced inward currents of similar amplitude (Figure 2a, b). In addition, BvPMT5a-expressing oocytes were challenged with the glucose derivative glucuronic acid, the hexose deoxy sugars fucose and rhamnose, the aldopentose arabinose, various linear polyols (sorbitol, mannitol, glycerol, xylitol) and the cyclic polyol myoinositol. Among these, mannitol, glucuronic acid and glycerol only produced very weak inward currents. Sorbitol, arabinose, fucose, rhamnose and myo-inositol induced similar currents to glucose and fructose, while xylitol caused the largest current response (Figure 2b). This behavior of BvPMT5a points to a transporter of broad substrate specificity. Due to the favorable signal-to-background-noise ratio with xylitol as a BvPMT5a substrate, this polyol was selected as a representative substrate to study the involvement of protons as a potential co-substrate in the translocation process. As expected from a H^+^-driven monosaccharide/polyol transporter, the polyol-induced inward currents became smaller in the presence of 10 mM xylitol when, at a membrane voltage of −40 mV, the external pH was increased from 5.5 to 6.5 and 7.5, leading to a gradual decrease in the proton motive force (PMF) (Figure S6a). A complete loss of the PMF was achieved by decreasing the membrane voltage to 0 mV and buffering the external pH to 7.5 (pH of the cytosol) (Figure S6b). Under this condition, xylitol application, however, still elicited inward currents that reached about 25% of those driven by a 100-fold H^+^ gradient at pH 5.5 (Figure S6b). In the absence of a PMF, H^+^ uptake into the cell was driven solely by the polyol concentration gradient directed into the cytosol. When the membrane voltage became increasingly hyperpolarized, the PMF increased, resulting in larger inward currents under any pH situation. However, at voltages more negative than −80 mV, the xylitol-induced currents measured at pH 6.5 and 5.5 became very similar, suggesting that the maximal transport capacity reached a similar level under both pH conditions and is no longer promoted by the voltage part of the PMF. When instead of the external pH, the xylitol concentration was varied by adding either 1, 3, 5, 10, 20, 30, 50 or 100 mM xylitol, the inward current increased stepwise from a concentration of 1 to 20 mM and saturated above 30 mM (Figure 3a, b). This saturation behavior could be fitted with a Michaelis-Menten function from which a K_m_ value of 2.5 mM was derived (Figure 3b, c). When the electrical driving force was increased from −40 to −120 mV, thus near to the resting membrane voltage of the taproot parenchyma cells (Figures 1b, S2b), the affinity to this polyol substrate increased almost two-fold as the K_m_ dropped from 2.5 to 1.5 mM (Figure 3c). In analogous experiments involving the glucose-dose dependency of BvPMT5a (Figure 3d-f), the derived K_m_ values for glucose also decreased (Figure 3e, f), indicating that the PMF is energizing the BvPMT5a H^+^/glucose cotransport. This identifies BvPMT5a as a potential candidate for glucose uploading in the beet taproot.

**Figure 2.**
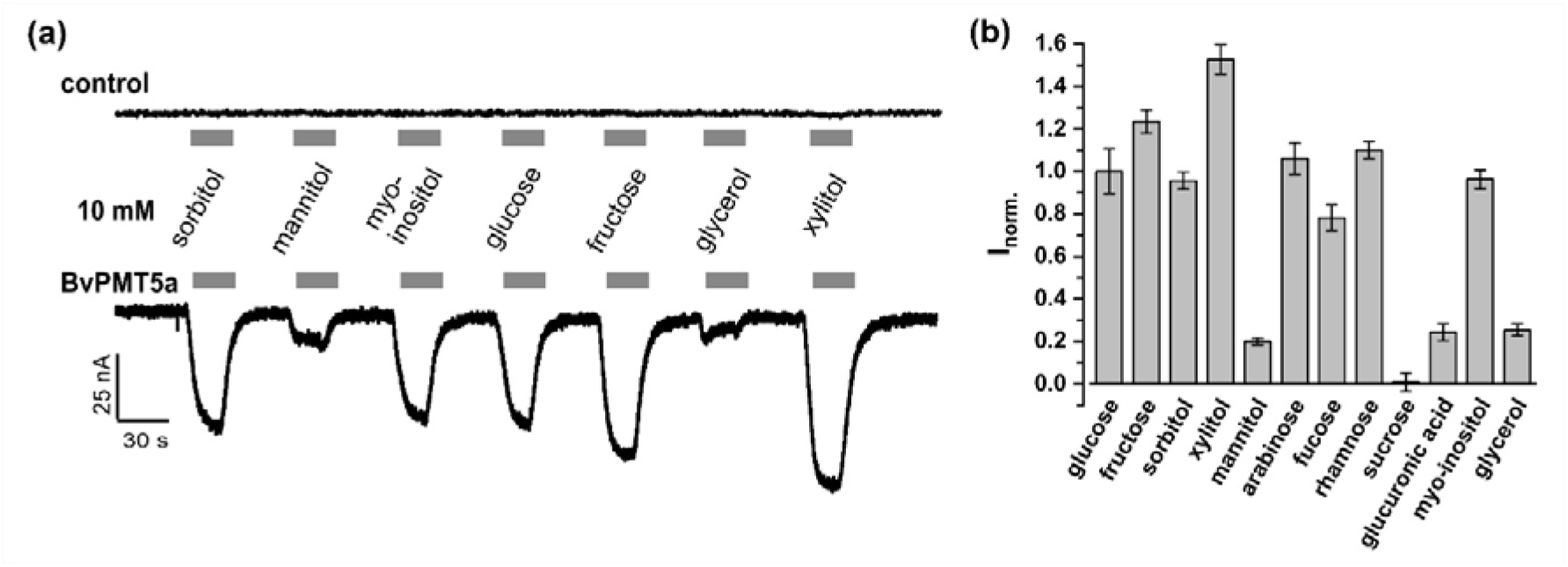
Substrate specificity of BvPMT5a. (a) Representative macroscopic current recordings of *Xenopus laevis* oocytes injected either with water (control) or BvPMT5a complementary RNA. Currents were recorded at a membrane potential of −40 mV and at pH 5.5 during a 30 s application (grey bar) of different sugar compounds (10 mM). Downward deflections indicate inward currents. (b) Current responses of BvPMT5a-expressing oocytes to the application of different sugar compounds (10 mM). The respective responses of each oocyte were normalized to the glucose-induced change in the inward currents of that oocyte. Data represents means ± SEM of 6 to 36 individual oocytes.

**Figure 3.**
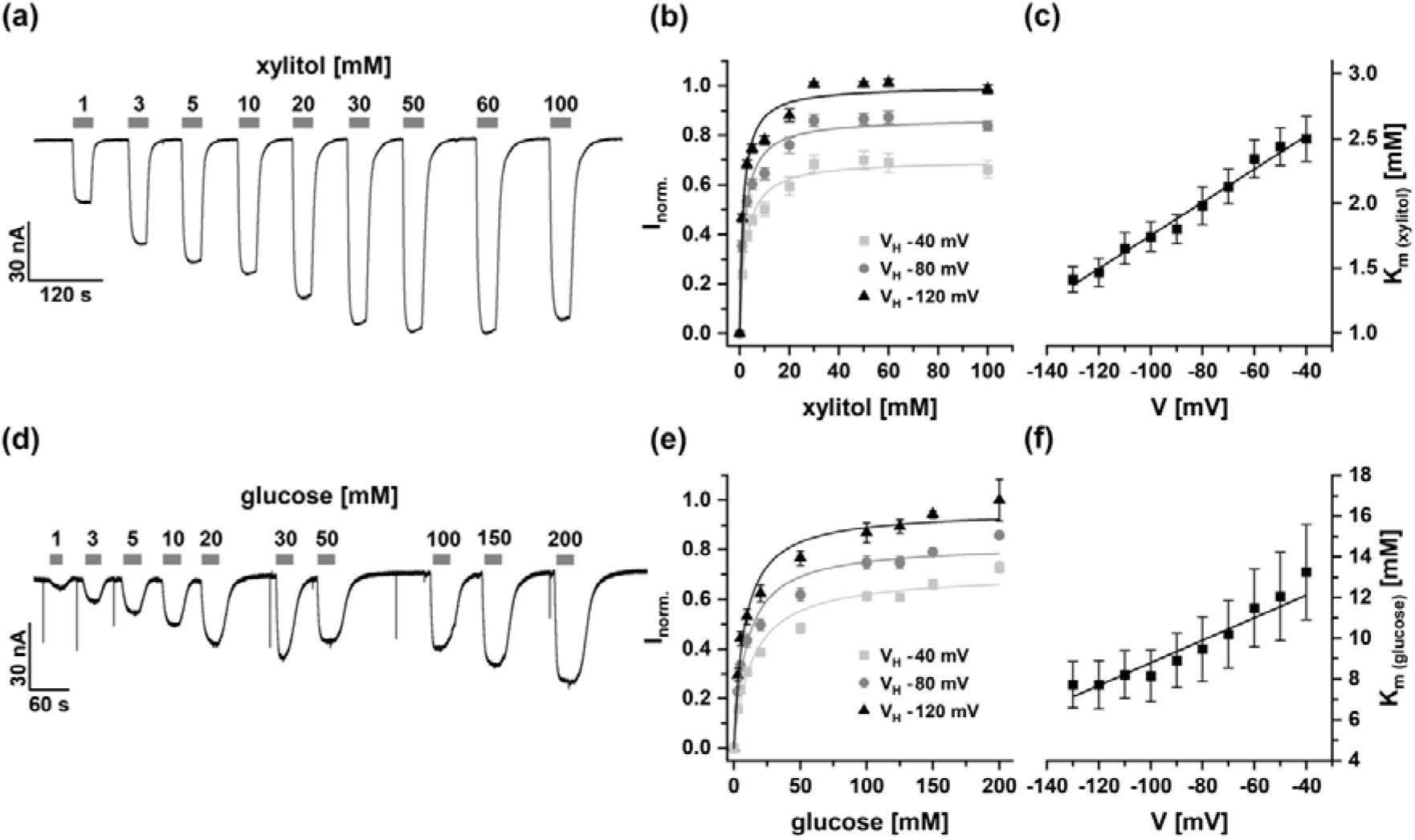
Xylitol- and glucose-dose dependency of BvPMT5a. (a, d) Current responses of BvPMT5a-expressing oocytes to xylitol (a) or glucose (d) application at the indicated concentrations. The duration of the substrate administration is indicated by the grey bar above the current trace. Currents were recorded at a membrane voltage of −40 mV at pH 5.5. (b, e) Xylitol- or-glucose-dependent currents plotted as a function of the substrate concentration. Currents were recorded at the membrane voltages as indicated and pH 5.5. Recorded currents were normalized to the maximum current recorded at a membrane voltage of −120 mV. The solid line gives the best fit of the data set with a Michaelis-Menten function. (c, f) Voltage dependency of the K_m_ values. K_m_ values derived from the best Michaelis-Menten-fits as shown in b, e were plotted against the respective membrane voltages. In b, c and e, f, data represents means ± SEM of 9 and 10 individual oocytes, respectively.

### BvSTP13 is a high-affinity, proton-coupled glucose and sucrose transporter

Like BvPMT5a, BvSTP13 was heterologously expressed in *Xenopus laevis* oocytes characterized its transport features. At an external pH of 5.5 and a membrane potential of −40 mV, BvSTP13-expressing oocytes were exposed to various monosaccharides as well as to di- and trisaccharides (Figure 4a). Upon application of 10 mM hexose quantities, BvSTP13-mediated inward currents of similar large amplitudes were recorded with glucose, fructose, galactose and mannose. In contrast, the polyols sorbitol, myo-inositol and xylitol, the hexose deoxy sugars fucose and rhamnose, the aldopentose arabinose and the hexose derivative glucuronic acid all caused no or only small currents. The aldopentose xylose, however, triggered current responses that reached approximately 70% of those obtained with glucose. When exposed to the glucose-fructose disaccharide sucrose, similar pronounced inward currents to those with xylose were obtained. Unexpectedly, even the trisaccharide raffinose evoked current responses of amplitudes that were still about 40% of those reached with glucose. BvSTP13 also accepted the fluorescent sucrose surrogate esculin as a substrate for the H^+^-coupled uptake into the oocytes (Figure S7a). The esculin transport capability of BvSTP13 was confirmed using tobacco mesophyll protoplasts from *N. benthamiana* leaves transiently transformed with BvSTP13. After incubation of control and BvSTP13 protoplasts in 0.5 mM esculin, fluorescence signals were detected only in BvSTP13-expressing protoplasts (Fig. S8). The fluorescent probe accumulated in the vacuoles (Fig. S8), suggesting that after BvSTP13-meditiated uptake of esculin from the extracellular space into the cytosol, the fluorescent probe was further shuttled into the vacuole via endogenous TST-like sucrose antiporter (Rottmann et al. 2018; Jung et al., 2015). The current responses from BvSTP13-expressing oocytes together with these esculin-loaded BvSTP13-equipped plant cells demonstrate that BvSTP13 is not a typical hexose transporter; it is capable of transporting not only certain monosaccharides but also sucrose and raffinose across the plasma membrane.

**Figure 4.**
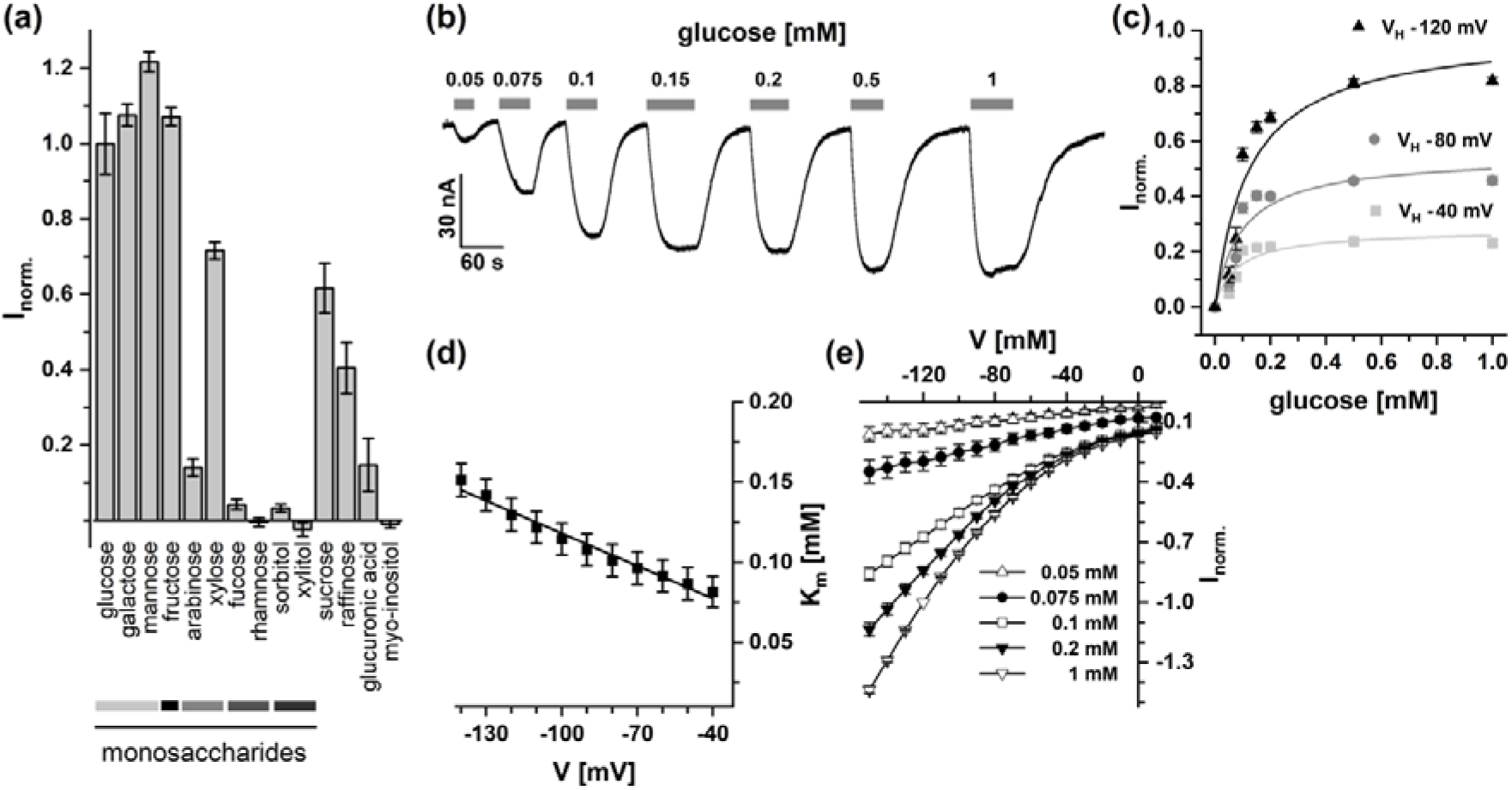
Substrate specificity, glucose-dose and voltage dependency of BvSTP13. (a) Current responses of BvSTP13-expressing oocytes to the application of different sugar compounds (10 mM) recorded at a membrane voltage of −40 mV. The respective responses of each oocyte were normalized to the glucose-induced change in the inward currents of that oocyte. Data represents means ± SEM of 7 to 24 individual oocytes. Grey scaled bars marked different groups of monosaccharides such as aldohexoses, ketohexose, aldopentoses, desoxypentoses and sugar alcohols (from left to right). (b) Representative current responses of BvSTP13-expressing oocytes to glucose application at the indicated concentrations. The duration of glucose administration is indicated by the grey bar above the current trace. Currents were recorded at a membrane voltage of −40 mV at pH 5.5. Noise peaks generated during the perfusion were dimmed offline. (c) Glucose-dependent BvSTP13-mediated currents plotted as a function of the substrate concentration. Currents were recorded at the membrane voltages indicated and pH 5.5. Recorded currents were normalized to the maximum current recorded at a membrane voltage of −120 mV. The solid line gives the best fit of the data set with a Michaelis-Menten function. (d) Voltage dependency of the K_m_ values. K_m_ values derived from the best Michaelis-Menten-fits as shown in C were plotted against the respective membrane voltages. (e) Current-voltage curves recorded at pH 5.5 under glucose treatment at indicated concentrations. Currents measured were normalized to the response to 1 mM glucose measured at −120 mV. In a-e all experiments were conducted at pH 5.5. In c-e, data represents means ± SEM of 12 individual oocytes.

To determine the glucose-dose dependency of the BvSTP13 transporter, the glucose concentration was increased stepwise from 0.05 mM to 1.0 mM (Figure 4b). In these experiments, inward currents were evoked with as little as 0.05 mM glucose. Currents tended to saturate when the substrate concentration was raised above 0.1 mM (Figure 4b, c). The K_m_ value at a membrane potential of −40 mV for glucose was 0.075 mM (Figure 4c, d), indicating that BvSTP13 represents a high-affinity sugar transporter. Like glucose, the application of sucrose elicited H^+^ inward currents at a concentration as low as 0.05 mM (Figure S7a, b), suggesting that BvSTP13 also has a high affinity to sucrose. At the glucose and sucrose concentrations tested, membrane hyperpolarization and an acidic external pH enhanced the inward currents (Figures 4e, S7b, S9). Together, the voltage and pH dependency of the BvSTP13-mediated currents demonstrate that as with BvPMT5a, the BvSTP13-mediated sugar translocation is proton-coupled, so thermodynamically driven by the proton motive force and the sugar gradient (cf. Carpaneto et al., 2005; Reinders et al., 2005; Wittek et al., 2017). However, in contrast to BvPMT5a (Figure 3f), the K_m_ values of BvSTP13 for glucose surprisingly increased to about 0.16 mM upon hyperpolarization to −140 mV. This indicates that BvPMT5a gains a higher sugar affinity when the membrane potential is depolarized.

BvPMT5a and BvSTP13 were noticed because they were transcriptionally up-regulated under cold in taproot tissue (Figure S5b, c). Thus, we asked whether and how carrier function is affected by lower temperatures. In the Xenopus oocyte system, the temperature was decreased from 35 to 5 °C in 10 °C steps, and the glucose-induced current responses were monitored (Figure 5). Current amplitudes with both transporters decreased with each cooling step. However, BvPMT5a-mediated currents could only be resolved when the temperature was raised above 5 °C. Above this temperature threshold, warming up the oocyte by 10 °C steps increased the transporter activity with a Q_10_ of about 4. In contrast, significant BvSTP13-related currents were recorded already at 5 °C and were characterized by a Q_10_ of about 2.

**Figure 5.**
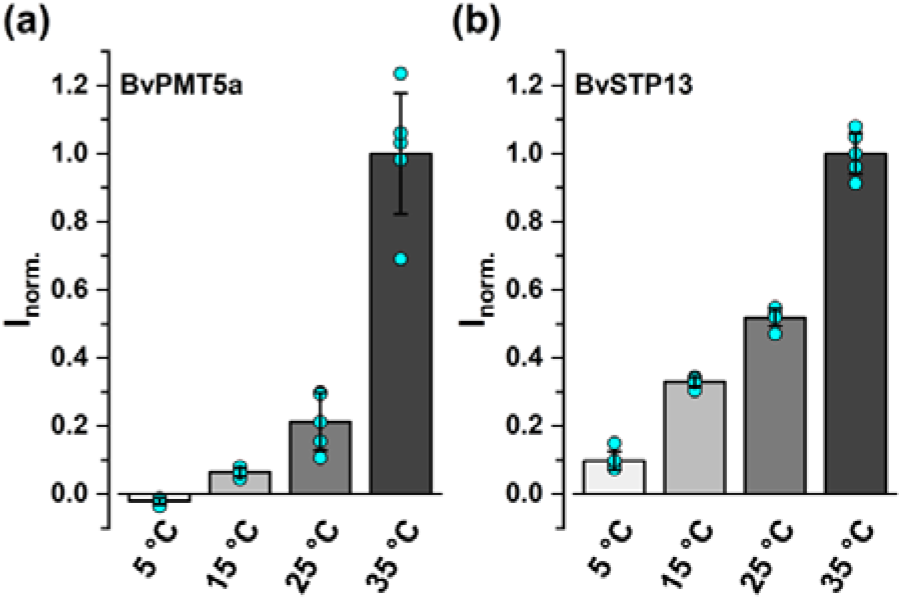
Temperature dependency of BvPMT5a and BvSTP13. Current responses of BvPMT5a (a)- and BvSTP13 (b)-expressing oocytes to 50 or 0.5 mM glucose, respectively, measured at −140 mV and temperatures as indicated were normalized to those at 35 °C. A Q_10_ value of 3.97 ± 0.57 for BvPMT5a and 2.18 ± 0.64 for BvSTP13 (means ± SD, n = 5) was determined. Q_10_ values were calculated as mean factors between temperatures, which resulted in distinct currents in the range of 15 - 35 °C for BvPMT15 and 5 - 35 °C for BvSTP13 (mean ± SD of 5 individual oocytes). Turquoise circles indicate the individual data points.

### Modeling the BvSTP13 structure with bound mono- and disaccharide

To obtain the first insights into the molecular nature of the broad sugar specificity of BvSTP13, we modeled BvSTP13, based on the known structure of the monosaccharide transporter AtSTP10 from *Arabidopsis thaliana* (Rottmann et al., 2016; Paulsen et al., 2019). In accordance with their shared overall 6TM-loop-6TM topology (TM, transmembrane domain), the BvSTP13 amino acid sequence could be perfectly mapped onto the AtSTP10 structure (Figure S10). The 3D model obtained for BvSTP13 revealed all the structural hallmarks of the STP protein family. This includes the existence of a characteristic ‘lid-domain’ covering the extracellular entry pathway to the sugar binding site, and a cavity that is formed between the N-terminal and C-terminal halves of the sugar transporter (Figure S10a). The structural alignment further revealed that core amino acid residues, identified as constituting the binding sites for coordinating the glucose substrate (Paulsen et al., 2019), were perfectly conserved between both transporters, apart from Leu43 in AtSTP10, which is replaced conservatively by valine (Val44) in BvSTP13 (Figure S10b). The presence of a hydrophilic polyethylene glycol (PEG) moiety above the bound glucose molecule in the AtSTP10 structure indicates that the saccharide binding cleft in the determined outward open conformation is wide enough to accommodate carbohydrates larger than monosaccharides (Figure 6). In our model of BvSTP13 bound with sucrose, space for the second carbohydrate moiety of the disaccharide is provided by changes in the sidechain conformation of Asn304 and Met307. This suggests that the spatial requirements of sucrose accommodation could seemingly be fulfilled in BvSTP13 as well as in AtSTP10.

**Figure 6.**
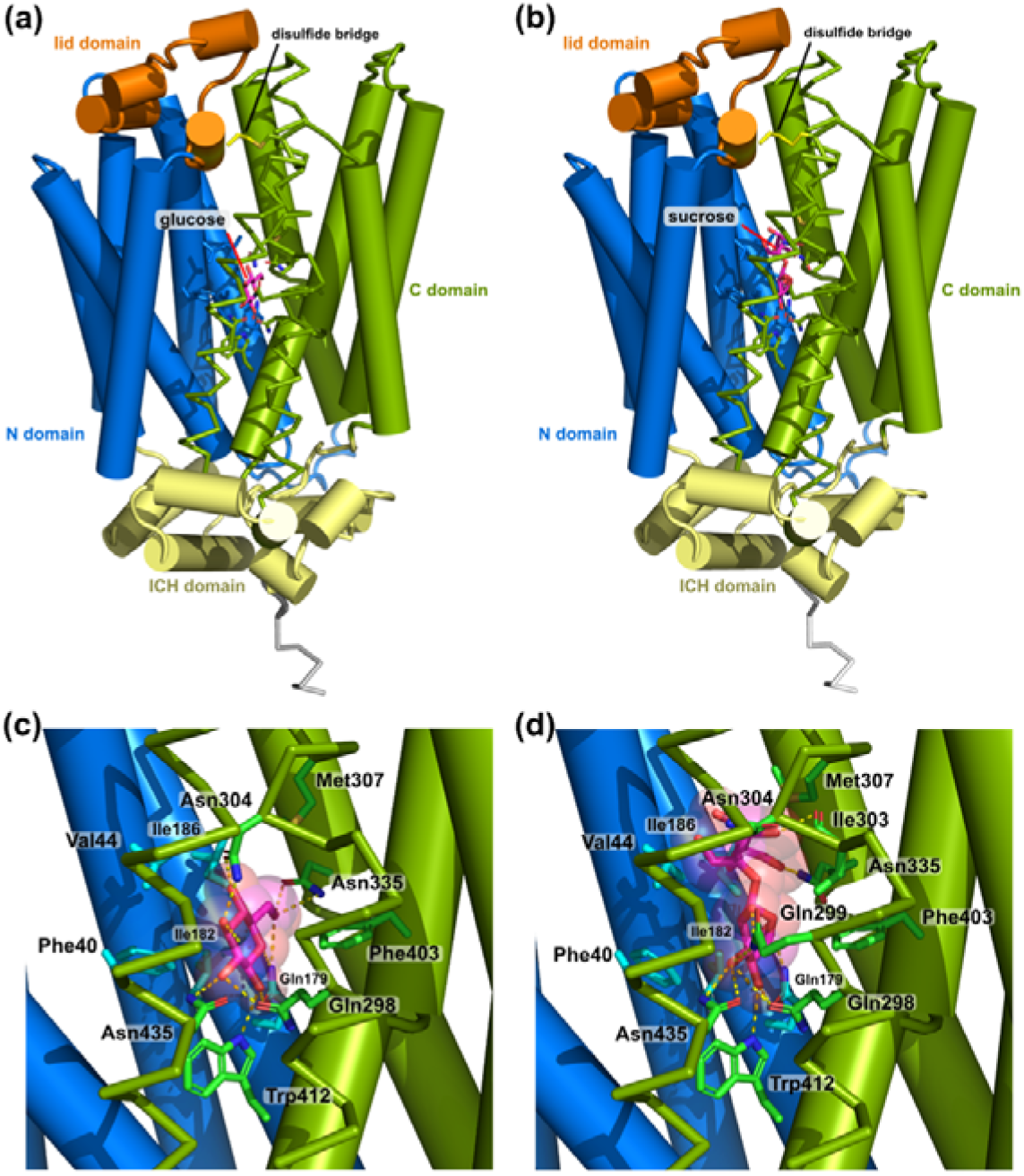
Comparison of a BvSTP13 model bound to glucose and sucrose. (a) A BvSTP13 model with glucose bound in the saccharide binding site. (b) BvSTP13 3D model bound to sucrose. The BvSTP13 model follows the color-coding scheme to indicate the different domains as stated in Figure S10. The saccharide moieties are shown as stick representation with their carbon atoms colored in magenta and the oxygen atoms marked in red. (c) An enlargement of the binding site in the interaction of BvSTP13 with glucose is shown. Residues in close proximity (≤ 5 Å) are shown as sticks, hydrogen bonds between the glucose molecule and residues of BvSTP13 are indicated by stippled lines colored in yellow. (d) The binding site of a BvSTP13 model interacting with sucrose is shown (magnification). The sucrose molecule was docked into the saccharide binding site of our BvSTP13 homology model such that the glucose moiety of the sucrose occupied the same position as the glucose molecule in the AtSTP10 structure. Energy minimization was performed, and a 10 ns MD trajectory was calculated with BvSTP13 placed in a bilayer membrane and the extra- and intracellular part surrounded by water. The model shows that the glucose moiety of the sucrose molecule can engage in similar hydrogen bonds as the glucose in the BvSTP13-glucose model. Furthermore, the fructose moiety of the sucrose molecule can form hydrogen bonds for instance with the carbonyl of Ile303 and with the carboxyamide group of Asn335.

The proton donor/acceptor residue pair needed for proton translocation, Asp42 and Arg142 in AtSTP10, is preserved in BvSTP13 as well as in the polyol transporter BvPMT5a. However, amino acid residues involved in substrate coordination partially differ between polyol and sugar transporters. Our modeling approach revealed that the ‘lid domain’ differs between AtSTP10 and BvSTP13, in particular, the non-structured loop C-terminal to the two short helices in the apoplastic loop between TM1 and TM2 (Figure S10a). Markedly, BvPMT5a lacks a ‘lid domain’. This ‘lid domain’ (Figure S11) is specific for members of the STP family. It covers the extracellular entry pathway of the substrate binding cavity. Via a conserved disulfide bond, the ‘lid domain’ links the N- and C-terminal halves of STP-type transporters, possibly allowing the propagation of structural rearrangements between both domains during the transport cycle.

## Discussion

### BvPMT5a and BvSTP13 together function as low and high-affinity proton-driven importers with broad substrate specificity

Sucrose in the plant is disseminated from source to sink via the phloem network. In sink organs such as a taproot in the case of sugar beet, sucrose exits the phloem apoplastically (Lemoine et al., 1988; Godt & Roitsch, 2006). However, depending on the activity of extracellular invertase at the exit site (Lemoine et al., 1988; Jammer et al., 2020), parenchyma sugar transporters will be faced with glucose and fructose in addition to sucrose. Our electrophysiological studies with taproot parenchyma cells clearly demonstrate that the plasma membrane responds to glucose and sucrose, as expected from transporter-mediated proton-driven sugar import (Figures 1, S2). Two taproot-expressed transporters, BvPMT5a and BvSTP13, have been identified in the oocyte system as H^+^/solute symporters with broad substrate specificity.

BvPMT5a mediates the proton-coupled uptake of glucose with a millimolar affinity (Figure 3). BvSTP13 shuttles both glucose and sucrose with submillimolar affinities (Figure 4, Figure 6a, b). Thus together, BvPMT5a and BvSTP13 provide low and high-affinity glucose uptake (Figures 3f, 4d). Whether the opposite weak voltage dependency of the BvSTP13 glucose affinity is correlated with a spatial rearrangement of substrate binding sites during the transport cycle conferred by the ‘Lid domain’ – present in BvSTP13 but absent in BvPMT5a (Figure S11) – needs to be explored in further studies.

### How to fit glucose and sucrose in the same transporter?

We used homology models for BvSTP13 and MdSTP13a (Li et al., 2020) and in silico docking of various saccharides to unravel how possible differences in saccharide binding in different members of the STP sugar transporter family might explain their observed substrate specificity (Figures 6, S10). In line with their similar saccharide specificity, all amino acid residues in the binding cleft of the STP13 sugar transporter from sugar beet and apple (Li et al., 2020) were conserved within an 8 Å sphere of the glucose moiety position as found in AtSTP10. Likewise, in this region only three amino acids differ between AtSTP10 and BvSTP13, all other residues are identical (see also Figure S10b). Based on the presence of three water molecules and a hydrophilic polyethylene glycol molecule in the binding cleft just above the glucose moiety, our MD simulations suggest that the binding site exhibits plasticity. This binding cleft plasticity could allow to accommodate di- and trisaccharides in BvSTP13 (and possibly AtSTP10). The three amino acids, that differ between AtSTP10 and BvSTP13 are located below a conserved tryptophan (Trp410 and Trp412 in AtSTP10 and BvSTP13, respectively). This residue resides beneath the saccharide binding cleft occupied by glucose and can be considered a kind of dead-end of the sugar accommodating cavity. The currently available structure (of only the outward open conformation) (Paulsen et al., 2019) can only provide insights into the binding situation when the saccharide moiety enters the cleft and is coordinated at the glucose binding site. Future work needs to elucidate whether the carbohydrate specificity of STP transporters is determined by the binding site itself or is attributable to conformational changes during their predicted outward open to inward open transport cycle.

### BvPMT5a and BvSTP13 with a possible role in response to low temperatures

As with the plasma membrane sugar transporters BvPMT5a and BvSTP13, the expression of the vacuolar Arabidopsis TONOPLAST SUGAR TRANSPORTERS TST1 and TST2 are induced by low temperatures (Wormit et al., 2006). During cold acclimation, *tst1/2* knockout lines exhibited elevated sucrose levels, but reduced glucose and fructose levels in the leaves compared with wild-type plants while showing a lower cold tolerance (Wormit et al., 2006; Klemens et al., 2014). Thus, the cellular sugar content contributes to cold hardening. These observations suggest that for cold tolerance of the sugar beet taproot, BvSTP13- and BvPMT5a-mediated plasma membrane hexose transport may be important. In addition, polyols and raffinose are also protective towards low temperatures (Wanner & Junttila, 1999; Gusta et al., 2004; Krasensky & Jonak, 2012; Klemens et al., 2014; Tarkowski & Van den Ende, 2015; Keller et al., 2021), and these are also substrates of BvPMT5a and BvSTP13, respectively. Therefore, one could assume that BvPMT5a and BvSTP13 could provide root parenchyma cells with cold protective compounds. Given that the temperature activity profiles of the two sugar transporters overlap, with BvSTP13 being more cold-tolerant than BvPMT5a, it is tempting to speculate that during cold acclimation, these H^+^ symporters work hand in hand. Generation of suitable sugar beet mutants, their characterization and *in vivo* transporter localization assays, are required to assess the physiological importance of BvPMT5a and BvSTP13 in beet taproot sugar uptake in general and cold tolerance in particular.

### Experimental procedures

#### Plant material and cultivation

*Beta vulgaris* plants (GT1-3 hybrids, Lisanna) were kindly provided by KWS SAAT SE (Germany). For electrophysiological characterization and RT-qPCR analysis, sugar beet plants were cultivated on Profi Substrat soil (CL ED73 Puls + Eisen, Einheitserde Werkverband e.V) under a 14/10 h day/night regime with a light intensity of about 120 μmol m^−2^ s^−1^ (sodium vapor lamp Sonte Agro 400). The temperature was 24 C and the relative humidity 60%. For transient transformation, *Nicotiana benthamiana* plants were grown in the green house under conditions described in Reyer et al. (2020).

#### Membrane voltage recordings in sugar beet taproots

Plants of 95 to 115 days old were used for membrane voltage recordings. The taproots were harvested, and cross sections of whole root prepared (middle part of the root, 0.5 mm slice). From these slices, the periderm was detached with sharp forceps to create a window. The tissue in this window was cut tangential to get a sample of 10 to 15 cell layers thick. The sample was glued with medical adhesive B liquid (ULRICH Swiss, St Gallen, Switzerland) to a cover glass, which was mounted with double-sided adhesive tape in the lid of a 3 cm Petri dish. It was bathed in 3 mL measuring solution (1 mM CaCl_2_, 1 mM KCl, 10 mM 2-(N-morpholino)ethanesulfonic acid (MES), adjusted with Bis-Tris propane (BTP) to pH 6.0) and the sample was incubated for 16 h at 18 °C in the dark. The solution was changed 1 h before measurement, and samples remained in the dark. The free-running membrane voltage recordings were essentially performed as described (Reyer et al., 2020). Briefly, glass microelectrodes made of borosilicate glass capillaries (length 100 mm, Ø_outer_ 1.0 mm, wall thickness 0.21 mm, Hilgenberg GmbH, Malsfeld, Germany) and filled with 300 mM KCl were impaled into the cells under microscopic control. The acquired data were analyzed using Microsoft Excel 2010 and Origin Pro-2021.

#### Proton flux measurements on sugar beet taproots

For application of the scanning ion-selective electrode (SISE) technique (Dindas et al., 2018), taproots were cut into slices (Ø ∼ approx. 0.5 cm) and an intact layer of parenchyma cells was dissected from a small area of such a slice. The entire slices were then immediately incubated in a basic salt medium (BSM, 0.5 mM KCl; 0.1 mM CaCl_2_, pH 5.3 unbuffered) and left overnight in the dark at room temperature. One hour prior to measurement, the slices were mounted with non-woven microporous adhesive tape (URGOPORE, 1.25 cm, Urgo Medical, Chenôve, France) on Petri dishes (diameter 8.5 cm) and subsequently filled with 25 mL of BSM. Thirty minutes prior to measurement, the BSM solutions in the petri dishes were changed. Net H^+^ fluxes were then measured from the exposed cells using non-invasive H^+^-selective scanning microelectrodes. According to (Newman, 2001) and (Dindas et al., 2018), microelectrodes were pulled from unfilamented borosilicate glass capillaries (Ø 1.0 mm, Science Products GmbH, Hofheim, Germany) dried over night at 220 °C, then silanized with N,N-dimethyltrimethylsilylamine (Sigma-Aldrich) for 1 h. The electrodes were subsequently back-filled with a backfilling solution (15 mM NaCl/40 mM KH_2_PO_4_, pH adjusted to 6.0 using NaOH for H^+^) and front-filled with an H^+^-selective ionophore cocktail (catalogue number 95291 for H^+^, Sigma-Aldrich). Calibration of H^+^-selective electrodes was performed at pH 4.0, pH 7.0 and pH 9.0. Electrodes with slope > 50 mV per decade and correlation > 0.999 were used for measurements. After calibration, the electrode was placed at a distance approximately 40 µM from the taproot sample using a SMC17 micromanipulator (Narishige Scientific Instrument Lab) and an upright microscope (Axioskop; Carl Zeiss AG, Oberkochen, Germany). During measurements electrodes were moved between two positions, i.e., close to and away from the sample (40 µm and 140 µm, respectively) at 10 s intervals using a microCstepping motor driver (US Digital, Vancouver, WA, USA). The difference in the potentials between these two points was recorded with a NI USB 6259 interface (National Instruments), controlled by a customCmade, LabviewCbased software ‘Ion Flux Monitor’. The recorded potential was converted offline into proton flux values using the LabviewCbased program ‘Ion Flux Analyser’, Microsoft Excel 2010 and Origin Pro 2021 software.

#### Esculin uptake in *Beta vulgaris* taproot cells

Taproot samples were prepared as described for membrane potential recordings. Thin slices were incubated in standard impalement solution with 0.5 mM esculin in measuring solution (1 mM CaCl_2_, 1 mM KCl, 10 mM 2-(N-morpholino)ethanesulfonic acid (MES), adjusted with Bis-Tris propane (BTP) to pH 6.0) at room temperature in the dark for 1, 5, 90 or 180 min followed by two washing steps (total duration, 15 min) on ice in 4 °C cold standard measuring solution without esculin. Afterwards the esculin fluorescence (emission filter 410 - 490 nm) was monitored with a confocal laser scanning microscope (Leica TCS SP5 II; Leica Mikrosysteme Vertrieb GmbH, Germany) using a 405 nm UV diode for excitation. Images were analyzed using the Leica confocal software and ImageJ (US National Institutes of Health; http://rsb.info.nih.gov/nih-image/).

#### Transient transformation and esculin loading of tobacco protoplast

*N. benthamiana* leaves were infiltrated with transgenic agrobacteria harboring pCambia2300 BvSTP13 as described elsewhere (Reyer et al., 2020). Three to four days after infiltration, mesophyll protoplasts were enzymatically isolated as follows. The transgene expressing leaf areas as well as untransformed control leaves were sandpapered (P2000-10µm grit) to remove the epidermis abaxial and placed in petri dishes (Ø 60 mm) with 3 ml enzyme solution (1 mM KCl, 1% (w/v) BSA (Albumin Fraction V), 0,05 % (w/v) pectolyase Y23 (Kyowa, Osaka, Japan), 0,5 % (w/v) cellulase R-10 (Yakult, Tokyo, Japan), 0,5 % (w/v) mazerozym R-10 (Yakult, Tokyo, Japan), 1 mM CaCl_2_, 10 mM MES/Tris, pH 5,6, adjusted with sorbitol to 480 mosmol kg^-1^ modified after Graus et al. (2018). The samples were incubated for 45-60 min with gentle horizontal shaking (30 rpm). After enzyme incubation, the extracted protoplasts were poured through a 100 µm mesh nylon net and washed with 20 ml of ice-cold protoplast buffer (1 mM CaCl_2_, 1 mM KCl, 10 mM 2-(N-morpholino)ethanesulfonic acid (MES), adjusted with Bis-Tris propane (BTP) to pH 6.0, adjusted to 480 mosmol kg^-1^ with sorbitol). The suspension was centrifuged for 7 min at 4 °C 60 x *g* with minimum acceleration and break. The liquid was then discarded, while the sedimented protoplasts were resuspended in residual buffer and gently mixed with esculin containing protoplast buffer to a final concentration of 0.5 mM esculin in 13 ml round-bottom tubes. After incubation at room temperature for 30-60 min, esculin was removed by centrifugation and an additional washing step with 5 ml of protoplast buffer. The protoplasts stored on ice were immediately imaged with the confocal laser scanning microscope.

#### Current recordings from *Xenopus laevis* oocytes

The two-electrode voltage-clamp technique was applied to *Xenopus laevis* oocytes injected with complementary RNA coding for BvPMT5a and BvSTP13 essentially as described by (Wittek et al., 2017). A standard bath solution was used for the membrane current recordings: 100 mM KCl, 1 mM CaCl_2_, 1 mM MgCl_2_, 1 mM LaCl_3_, adjusted to 220 mosmol kg^-1^ with D-sorbitol or sucrose. Solutions were adjusted either to pH 5.5 with MES/Tris buffer or to pH 6.5 and pH 7.5 with HEPES/Tris buffer. The following sugar compounds were added to the solutions at the concentrations indicated in the figure legends: L-(+)-arabinose, D-(–)-fructose, L-(-)-fucose, D-(+)-galactose, D-(+)-glucose, glycerol, D-glucuronic acid, D-mannitol, D-(+)-mannose, myo-inositol, D-(+)-raffinose, L-rhamnose, D-sorbitol, sucrose, xylitol. Sugar-induced current responses were determined by subtracting the current responses at the end of sugar application from those before sugar administration. For this, usually 150 ms lasting voltage pulses in the range of 0 to −140 mV were applied in 10-mV decrements before and during sugar application, starting from a holding voltage of −40 mV. Current responses to sugar application were also determined from continuous recordings at a constant membrane voltage. For substrate specificity, the related current response of each oocyte was normalized to the respective measured glucose-induced current and the standard errors of the normalized current responses from different oocytes were calculated. To also reflect the variability of the glucose-induced currents, the standard error was calculated from the absolute glucose-triggered currents of all measured oocytes and then normalized to the averaged glucose-induced current.

To study the temperature dependency of the sugar transporters in oocytes, the bath solution passed through a heat exchanger in contact with peltier elements on which the recording chamber was mounted. The temperature was measured with a small thermistor close to the oocyte. In these experiments, the bath solution for BvPMT5-expressing oocytes contained 75 mM NaCl, 50 mM sucrose, 1 mM CaCl_2_, 1 mM MgCl_2_, 1 mM LaCl_3_ and 10 mM MES/TRIS, pH 4.5. To induce uptake currents, sucrose was replaced by 50 mM D-(+)-glucose. The bath solution for BvSTP13-expressing oocytes contained 96 mM NaCl, 2 mM KCl, 1 mM CaCl_2_, 1 mM MgCl_2_, 1 mM LaCl_3_, 0.5 mM D-sorbitol, and 10 mM Mes/Tris, pH 4.5. To induce uptake currents, D-sorbitol was replaced by 0.5 mM D-(+)-glucose.

Data acquisition and offline analysis were performed using the software programs Patch Master (HEKA Electronik, Lambrecht, Germany), Microsoft Excel, Origin2021 (OriginLab Corporation, Northampton, MA 01060 USA) and IgorPro (Wave Metrics Inc., Lake Oswego, OR, USA).

#### Molecular cloning of sugar beet transporter

To generate constructs for heterologous expression of fluorophore-labelled or untagged *BvPMT5a* or *BvSTP13*, the corresponding coding sequence was amplified from cDNA of root tissue from cold-treated sugar beets. PCR-amplification of target sequences using the primer pNBI-BvPMT5-f (5’-GGG CTG AGG CTT AAT ATG AGT GAA GGA ACT AAT AAA GCC ATG −3’) together with BvPMT5-pNBI16/21-r (5’-ATT CGC TGA GGT TTA GTG ATT GTC ATT TGT AAC AGT AGT ACT A −3’), or pNBI-BvSTP13-f (5’-ATT CGC TGA GGT TTA GTG ATT GTC ATT TGT AAC AGT AGT ACT A −3’) together with BvSTP13-pNBI16/21-r (5’-ATT CGC TGA GGT TTA TAG AGC TGC AGC TGC AGC AGA CCC ATT AT −3’) yielded PCR-fragments for cloning into pNBI16 (no tag), or pNBI21 (N-terminal fluorophore), respectively. PCR using the primer pairs pNBI-BvPMT5-f and BvPMT5-pNBI22-r (5’-CCA GGC TGA GGT TTA AGT GAT TGT CAT TTG TAA CAG TAG TAC TA −3’), or pNBI-BvHT2-f BvSTP13-pNBI22-r and BvSTP13-pNBI22-r (5’-CCA GGC TGA GGT TTA ATA GAG CTG CAG CAG ACC CAT TAT −3’), removed the stop codon of the transporter CDSs to allow generation of fusions to the N-terminus of the yellow fluorescent Venus (pNBI22), respectively. PCR fragments were directly cloned into the *PacI*-linearized expression vectors (pNBI16, pNBI21, or pNBI22) using the In-Fusion® HD Cloning Kit (Takara Bio USA, Inc.). Insert sequences were verified by sequencing (Eurofins, Germany).

#### Gene expression analysis

RNA extracted and sequenced in Martins Rodrigues et al. (2020) and Keller et al. (2021) was analyzed. For RT-qPCR analysis, sliced taproot tissues were frozen in liquid nitrogen, ground to a fine powder, and RNA extracted using the NucleoSpin(R) RNA Plant Kit (Machery-Nagel, Düren, Germany) according to the manufacturer’s guidelines. RNA transcription into cDNA was performed using the qScript cDNA Synthesis Kit (Quantbio, Beverly, USA). Expression analysis of *PMT5a* was conducted using the primers BvPMT5+439f (5’-ATC GCA CCT GTT TAC ACT GC −3’) and BvPMT5+656r (5’-GAC TCA GGC ATA GCA AGC AC −3’). These primers amplified the 217 bp-long *PMT5a* fragment at the chosen annealing temperature of 58 °C with an efficiency of 93%. Expression analysis of *STP13* was conducted using the primers BvSTP13+1031f (5’-ACT CTG TCG ACA AGC TTG GA −3’) and BvSTP13+1218r (5’-AGA CCA CGC AAA AGA GGA GA −3’). These primers amplified the 193 bp-long *STP13* fragment at the chosen annealing temperature of 58 °C with an efficiency of 88%.

#### Phylogenetics of sugar transporters

The phylogenetic relationships *Arabidopsis thaliana* and *Beta vulgaris* PMT and STP transporters were studied by aligning the derived amino acid sequences using the MUSCLE plugin (Edgar, 2004) within Geneious (Biomatters, Inc., San Diego, CA) with default parameters. The alignments were trimmed using trimAl v1.2rev59 (Capella-Gutierrez et al., 2009) using the implemented ‘gappyout’ algorithm. Phylogenetic reconstruction of the MSA was conducted using IQ-TREE multicore version 2.1.2 (Minh et al., 2020). The best-fit substitution models were LG+G4 and LG+I+G4 for the PMT and STP datasets, respectively, and were selected based on the Bayesian Information Criterion and implemented in the Maximum Likelyhood (ML) tree reconstruction. Branch support was estimated using 1000 replicates and ultrafast bootstrap (Hoang et al., 2018). Consensus trees were finally visualized employing the iTol online tool (https://itol.embl.de/).

#### 3D modeling of BvSTP13 and molecular dynamic analysis of saccharide binding

A 3D homology model of BvSTP13 was obtained based on the crystal structure of *Arabidopsis thaliana* STP10 (PDB entry 6H7D; Paulsen et al., 2019) using the modeling macro hm_build.mcr of the software package YASARA Structure version 20.12.24 (https://www.yasara.org, Krieger and Vriend, 2014; PMID 24996895). Briefly, the amino acid sequence of BvSTP13 covering residues Met1 to Leu537 was aligned to the sequence of STP10 using a PSI-BLAST search against the RCSB databank with a maximum E-value of 0.1 for template consideration. Three potential templates were identified: PDB entries 6H7D (AtSTP10, Paulsen et al., 2019), 4ZW9 (HsGTR3, Deng et al., 2015) and 5C65 (HsGTR3/SLC2A3; Pike A.C.W. et al. unpublished), although the latter two exhibited considerably lower scores in YASARA’s PSI-Blast search and alignment. For modeling a secondary structure prediction, a target sequence profile was built against related UniRef90 sequences. Fourteen initial homology models were then built using the template AtSTP10 (PDB entry 6H7D), employing five slightly different sequence alignments that differed in the adjustments of loop regions. Of these, two exhibited the best overall Z-scores of −0.432 and −0.465 in the YASARA scoring routine after energy minimization and molecular dynamics refinement. The model used for analysis comprised 505 amino acid residues harboring Gly12 to Ala516, the 11 N-terminal and the 21 C-terminal amino acids were not modeled because there is no template structure available for these residues. However, from the model it can be assumed that these residues are flexible and adopt a dynamic structure. The final model also contained the glucose moiety present in the original template AtSTP10 (PDB entry 6H7D), which engaged in an identical hydrogen bond pattern with surrounding residues in BvSTP13. This was because all amino acids in close proximity to the glucose moiety are conserved between AtSTP10 and BvSTP13. To obtain further insights into possible saccharide binding and specificity of BvSTP13, the monosaccharide fructose and the disaccharide sucrose were also docked in the saccharide binding site of the model. A short molecular dynamic simulation of the BvSTP13 model placed in a membrane layer with either bound glucose, fructose or sucrose was run using YASARA’s macro md_runmembrane.mcr. The hexaoxaicosandiol/PEG moiety that was part of the crystallization solution/condition of the original AtSTP10 crystal and which artificially occupied part of the inner binding cleft partially filled with the glucose molecule was removed before the MD simulation. The membrane region of the BvSTP13 molecule was predicted by YASARA. The BvSTP13 model was then centered in a box with the dimensions 83 x 83 x 113 Å and with the membrane region of the BvSTP13 model placed in the lipid bilayer comprising 159 phosphatidyl-ethanolamine molecules. Water was put above and below the membrane bilayer, sodium and chloride ions were placed at a concentration of 150 mM and to neutralize the protein charges within the box. After energy minimization of the membrane bilayer, the water solvent molecules and ions, an unrestrained molecular dynamic simulation was performed at 298K (25 °C) and constant pressure for 5 (BvSTP13 with glucose bound) and 10 ns (BvSTP13 with sucrose or fructose bound). The MD trajectories were analyzed with YASARA with respect to hydrogen bonding between the saccharide molecules and the sugar transporter.

#### Accession numbers

RNA-seq data are found in the GenBank Sequence Read Archive (BioProject PRJNA602804). The following accession numbers are listed in GenBank and RefBeet1.2: AtPMT1 (At2g16120), AtPMT2 (At2g16130), AtPMT3 (At2g18480), AtPMT4 (At2g20780), AtPMT5 (At3g18830), AtPMT6 (At4g36670), BvPMT3 (Bv6_137580_wnrf.t1), BvPMT4a (Bv8_183690_topt.t1), BvPMT4b (Bv9_206800_eujd.t1), BvPMT5a (Bv9_217740_uajm.t1), BvPMT5b (Bv8_196360_kwft.t1). AtSTP1 (At1g11260), AtSTP2 (At1g07340), AtSTP3 (At5g61520), AtSTP4 (At3g19930), AtSTP5 (At1g34580), AtSTP6 (At3g05960), AtSTP7 (At4g02050), AtSTP8 (At5g26250), AtSTP9 (At1g50310), AtSTP10 (At3g19940), AtSTP11 (At5g23270), AtSTP12 (At4g21480), AtSTP13 (At5g26340), AtSTP14 (At1g77210), BvSTP1a (Bv4_095750_qknz.t1), BvSTP1b (Bv8_181530_fpga.t1), BvSTP1c (Bv9_216930_ujxp.t1), BvSTP1d (Bv5_101120_supk.t1), BvSTP3a (Bv5_112890_mydf.t2), BvSTP3b (Bv5_120400_jtny.t2), BvSTP5 (Bv1_009250_xmsm.t1), BvSTP7 (Bv8_192620_dads.t1), BvSTP8 (Bv_004260_grqa.t1), BvSTP10a (Bv_006770_wdje.t1), BvSTP10b (Bv9_202750_dgpt.t1), BvSTP13 (Bv_008680_mhfn.t1), BvSTP14 (Bv8_197090_cxcd.t1). The accession numbers 6H7D (AtSTP10), 4ZW9 (HsGTR3) and 5C65 (HsGTR3/SLC2A3) refer to Protein Data Bank (PDB) entries.

## Acknowledgements

We thank Fábio Luiz Rogé Ferreira for his help in collecting raw electrophysiological data in Xenopus oocytes and Christina Müdsam for cloning of the sugar beet transporters. We are also grateful to Tracey Ann Cuin for critically proofreading this paper.

## Author contributions

R.H., I.M., H.E.N. designed the research; A.R., J.J., N.B., N.S., S.S. performed research; A.R., D.J., J.J., N.B., N.S., T.D.M., D.B., S.S., B.P. analyzed data and R.H., I.M, D.B., T.D.M., H.E.N. wrote the paper.

