## Supporting Information for "Beet taproot plasma membrane sugar transport revisited"

The following Supporting Information is available for this article:

**Figure S1.** Cellular loading of esculin in taproot cells.
**Figure S2.** Sucrose-induced changes in $H^+$ fluxes and membrane depolarization of *Beta vulgaris* taproot cells.
**Figure S3.** No mannitol-induced change in the membrane potential of *Beta vulgaris* taproot cells.
**Figure S4.** Phylogenetic tree of PMTs and STPs of *Arabidopsis thaliana* (At) and *Beta vulgaris* (Bt).
**Figure S5.** Expression of *SUT*-, *STP*- and *PMT*-like genes in *Beta vulgaris* taproots at ambient and low temperatures.
**Figure S6.** pH and voltage dependency of BvPMT5a.
**Figure S7.** Sucrose-dose dependency and voltage dependency of BvSTP13.
**Figure S8.** Loading of esculin in BvSTP13-transformed tobacco mesophyll protoplasts.
**Figure S9.** pH and voltage dependency of BvSTP13.
**Figure S10.** Comparison of the 3D homology model of BvSTP13 model and the crystal structure of AtSTP10.
**Figure S11.** Structure guided alignment of sugar transporters.

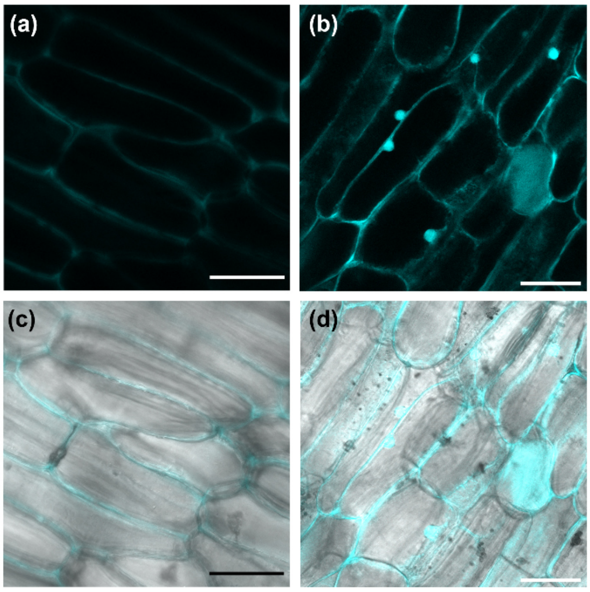


**Figure S1.** Cellular loading of esculin in taproot cells.

Fluorescent images of esculin fluorescence (upper row) and overlay (lower row) of fluorescent images with bright field of taproot slices from GT2 after incubation with 0.5 mM esculin for 1 min (a, c, z-stacks of 6 images), 180 min (b, d, single image) after washing. Scale bar: 40 µm. Number of biological replicates (= slices of different taproots) was at least n = 3.

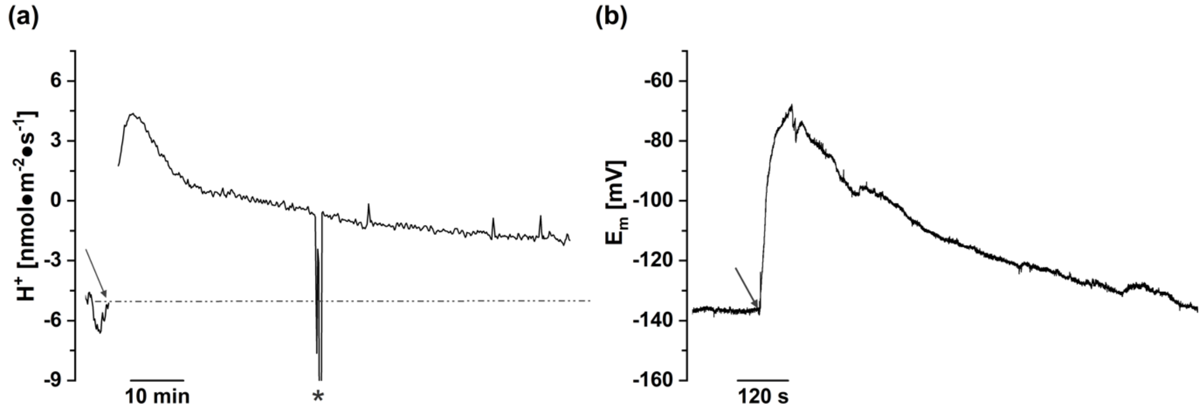


**Figure S2.** Sucrose-induced changes in $H^+$ fluxes and membrane depolarization of *Beta vulgaris* taproot cells. $H^+$ fluxes (a) and free running membrane voltage (b) recorded from a taproot slice in response to 50 mM sucrose treatment. Time of sucrose application is denoted by the arrow. Electrical disturbance indicated by (*). Negative and positive fluxes represent $H^+$ release from cells and $H^+$ uptake into cells, respectively. The $H^+$ flux level determined at rest shortly before sugar administration is indicated by a dotted line. In a, b, taproots from GT2 were used.

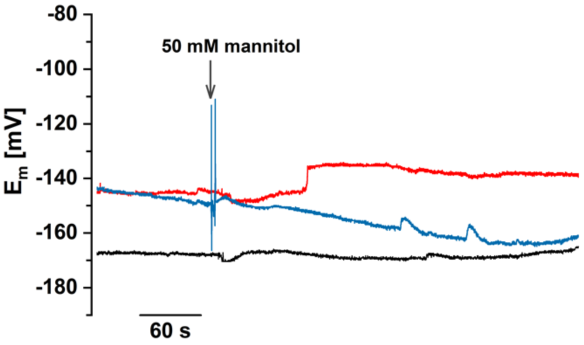


**Figure S3.** No mannitol-induced change in the membrane potential of *Beta vulgaris* taproot cells.

Recording of free running membrane potential from three different taproot slices (here color-coded) in response to 50 mM mannitol. The arrow indicates the time of mannitol application. The biological replicates were from the variety GT2.

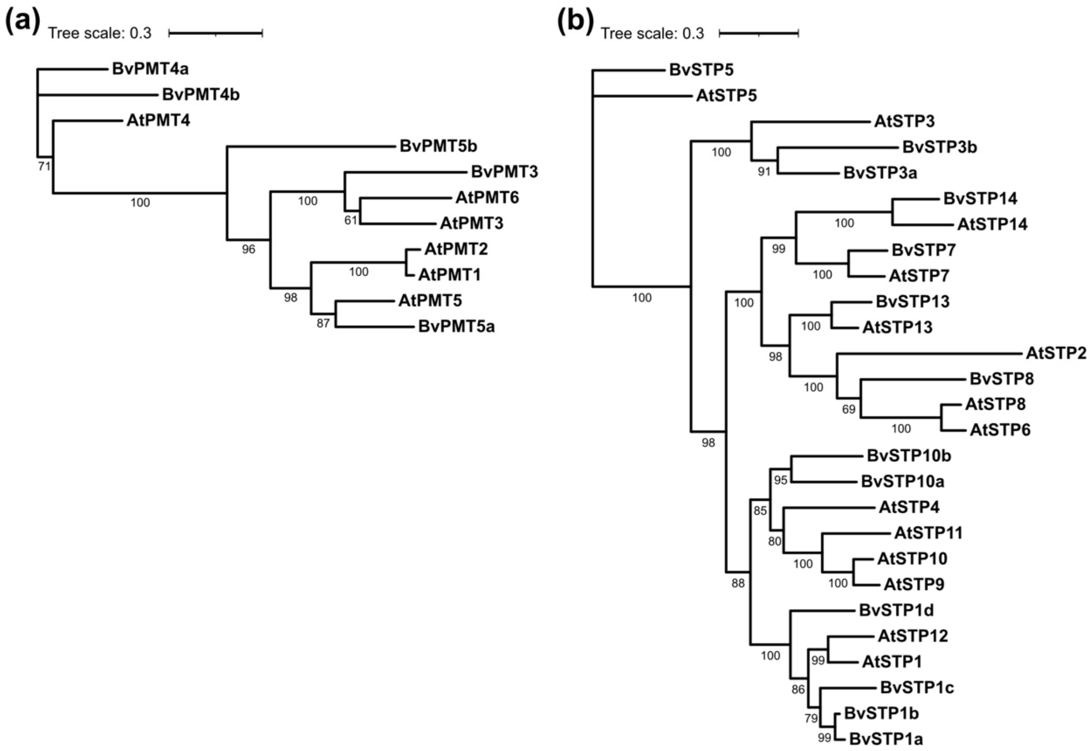


**Figure S4.** Phylogenetic tree of PMTs and STPs of *Arabidopsis thaliana* (At) and *Beta vulgaris* (Bt).

Consensus tree of the putative PMT (a) and STP proteins (b) of the two species. The accession numbers of the *Beta vulgaris* and *Arabidopsis thaliana* genes used are listed in RefBeet 1.2 and GenBank, respectively, as follows. In (a): AtPMT1 (At2g16120), AtPMT2 (At2g16130), AtPMT3 (At2g18480), AtPMT4 (At2g20780), AtPMT5 (At3g18830), AtPMT6 (At4g36670), BvPMT3 (Bv6_137580_wnrf.t1), BvPMT4a (Bv8_183690_topt.t1), BvPMT4b (Bv9_206800_eujd.t1), BvPMT5a (Bv9_217740_uajm.t1), BvPMT5b (Bv8_196360_kwft.t1). In (b): AtSTP1 (At1g11260), AtSTP2 (At1g07340), AtSTP3 (At5g61520), AtSTP4 (At3g19930), AtSTP5 (At1g34580), AtSTP6 (At3g05960), AtSTP7 (At4g02050), AtSTP8 (At5g26250), AtSTP9 (At1g50310), AtSTP10 (At3g19940), AtSTP11 (At5g23270), AtSTP12 (At4g21480), AtSTP13 (At5g26340), AtSTP14 (At1g77210), BvSTP1a (Bv4_095750_qknz.t1), BvSTP1b (Bv8_181530_fpga.t1), BvSTP1c (Bv9_216930_ujxp.t1), BvSTP1d (Bv5_101120_supk.t1), BvSTP3a (Bv5_112890_mydf.t2), BvSTP3b (Bv5_120400_jtny.t2), BvSTP5 (Bv1_009250_xmsm.t1), BvSTP7 (Bv8_192620_dads.t1), BvSTP8 (Bv_004260_grqa.t1), BvSTP10a (Bv_006770_wdje.t1), BvSTP10b (Bv9_202750_dgpt.t1), BvSTP13 (Bv_008680_mhfn.t1), BvSTP14 (Bv8_197090_cxcd.t1).

(a)

| famliy | name | gene ID | 20°C GT1 | 20°C GT2 | 20°C GT3 | 4°C GT1 | 4°C GT2 | 4°C GT3 | 0°C GT1 | 0°C GT2 | 0°C GT3 | putative location |
|---|---|---|---|---|---|---|---|---|---|---|---|---|
| SUT-like | BvSUT1 | gzum | 6066 | 3140 | 4956 | 4865 | 3552 | 5554 | 6348 | 5549 | 5144 | PM |
| | BvSUT3 | yemu | 2178 | 2144 | 2159 | 2124 | 1982 | 2208 | 2331 | 2991 | 2320 | PM |
| | BvSUT4 | zpft | 2390 | 2503 | 2163 | 5583 | 6019 | 4823 | 5442 | 5003 | 5314 | T |

(b)

| MST subfamliy | name | gene ID | 20°C GT1 | 20°C GT2 | 20°C GT3 | 4°C GT1 | 4°C GT2 | 4°C GT3 | 0°C GT1 | 0°C GT2 | 0°C GT3 | putative location |
|---|---|---|---|---|---|---|---|---|---|---|---|---|
| STP-like | BvSTP1a | qknz | 3 | 3 | 2 | 3 | 3 | 2 | 1 | 6 | 5 | PM |
| | BvSTP1b | fgpa | 1 | 1 | 1 | 1 | 1 | 1 | 1 | 1 | 1 | PM |
| | BvSTP1c | ujxp | 1 | 1 | 1 | 4 | 1 | 1 | 4 | 0 | 0 | PM |
| | BvSTP1d | supk | 1 | 1 | 7 | 3 | 11 | 8 | 96 | 129 | 7 | PM |
| | BvSTP3a | mydf | 16 | 10 | 18 | 73 | 74 | 120 | 173 | 84 | 182 | PM |
| | BvSTP3b | jtny | 3105 | 2522 | 2674 | 2838 | 2102 | 2053 | 2060 | 3039 | 1625 | PM |
| | BvSTP5 | xmsm | 223 | 56 | 132 | 56 | 41 | 178 | 76 | 28 | 60 | PM |
| | BvSTP7 | dads | 749 | 822 | 663 | 685 | 736 | 767 | 691 | 618 | 703 | PM |
| | BvSTP8 | grqa | 223 | 56 | 60 | 56 | 41 | 132 | 76 | 28 | 178 | PM |
| | BvSTP10a | wdje | 1 | 1 | 3 | 2 | 0 | 2 | 9 | 0 | 0 | PM |
| | BvSTP10b | dgpt | 1 | 1 | 1 | 0 | 0 | 0 | 1 | 0 | 0 | PM |
| | **BvSTP13** | mhfn | 403 | 71 | 179 | 1089 | 895 | 2767 | 11125 | 12797 | 7469 | PM |
| | BvSTP14 | cxcd | 391 | 340 | 265 | 947 | 2057 | 1745 | 5779 | 8917 | 5013 | PM |
| PMT-like | BvPMT3 | wnrf | 10 | 18 | 39 | 21 | 53 | 24 | 91 | 140 | 16 | PM |
| | BvPMT4a | topt | 692 | 747 | 640 | 655 | 470 | 708 | 793 | 668 | 522 | C/T |
| | BvPMT4b | eujd | 183 | 120 | 226 | 231 | 258 | 232 | 237 | 278 | 217 | C/T |
| | **BvPMT5a** | uajm | 315 | 218 | 217 | 1233 | 874 | 1534 | 3338 | 3045 | 3328 | PM |
| | BvPMT5b | kwft | 168 | 214 | 269 | 198 | 369 | 263 | 177 | 437 | 149 | PM |

low — high expression

(c)

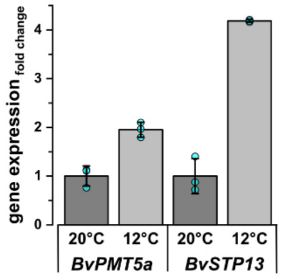


**Figure S5.** Expression of *SUT*-, *STP*- and *PMT*-like genes in *Beta vulgaris* taproots at ambient and low temperatures.

(a, b) Heat map presentation based on RNA-seq-sequence reads (cpm) of three different sugar beet hybrid genotypes (GT1 to GT3). Gene expression intensities were normalized to the highest and lowest value within each (sub)family. Numbers show RNA-seq sequence reads as means from three independent RNA-seg analyses per genotype and growth condition. The putative localization of the transporter is indicated as follows: plasma membrane (PM), plastid envelope (C), tonoplast (T). Plants were grown under control conditions (20 °C) for 10 weeks, transferred to 12 °C for one week, then grown for two more weeks at 4 °C, and finally for one week at 0 °C. In (a, b) the accession numbers of the *Beta vulgaris* genes used are listed in RefBeet 1.2 as follows. In (a) BvSUT1 (Bv1_000710_gzum.t1), BvSUT3

(Bv6_154300_yemu.t1), BvSUT4 (Bv5_124860_zpft.t1). Details of those gene identifiers given in (b) are found in Figure S4.

(c) Fold-change of *BvPMT5a* and *BvSTP13* expression in sugar beet root tissue before and after transfer of plants to low temperature. Plants from genotype "Lisanna" were grown for eight weeks at 20 °C, then one week at 12 °C. Samples were harvested at the corresponding temperatures at the indicated time points, and expression was quantified relative to *BvUGD1* expression. Relative expression at 20 °C was set to one. Bars represent means from n = 3 biological replicates ± SD. One replicate represents a pool of peeled taproots from at least three plants. Turquoise circles indicate the individual data points.

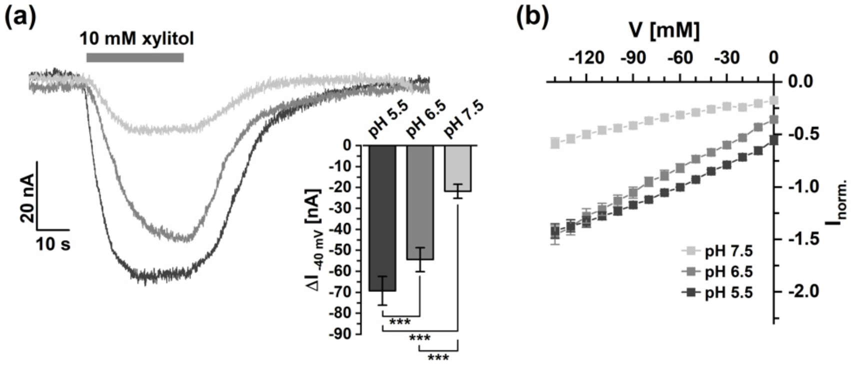


**Figure S6.** pH and voltage dependency of BvPMT5a.

(a) Left panel: Representative current traces recorded from a BvPMT5a-expressing oocyte at the indicated pH values and a membrane potential of -40 mV. Xylitol application (10 mM) is marked by a grey bar on top of the current trace. Downward deflections indicate inward currents. Right panel: Xylitol-induced current responses of different BvPMT5a-expressing oocytes quantified for the experimental conditions shown in the left panel. Data were analyzed for significant differences with paired Students *t*-test (***$p < 0.001$).

(b) Current-voltage curves recorded at pH 5.5, 6.5 and 7.5 under 10 mM xylitol treatment. BvPMT5a-mediated currents were normalized to the current response of that oocyte at pH 5.5 and -60 mV. Data in a (right panel) and b represent means ± SEM of 13 individual oocytes.

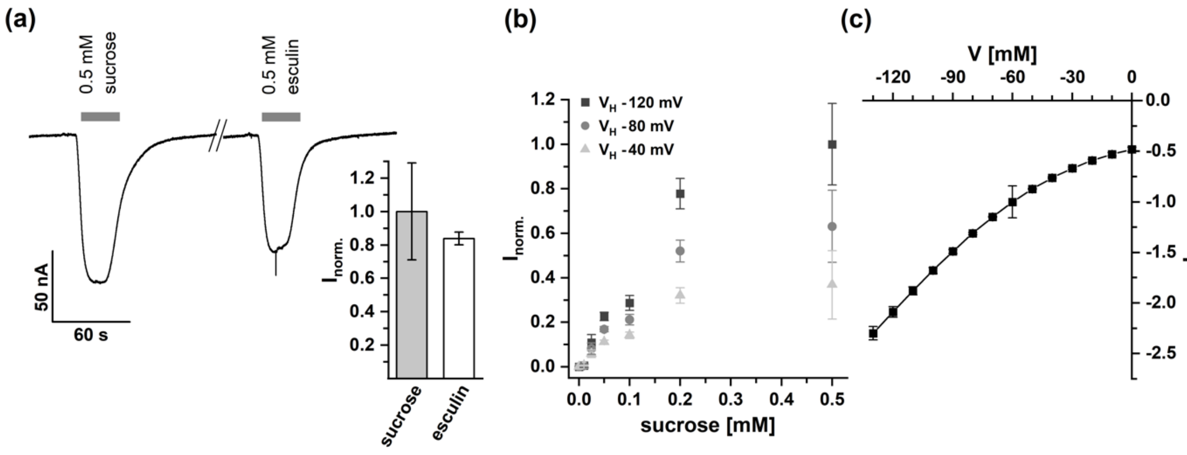


**Figure S7.** Sucrose-dose dependency and voltage dependency of BvSTP13.

(a) Current responses of BvSTP13-expressing oocytes to sucrose and esculin application at the indicated concentration. The duration of substrate administration is indicated by the grey bar above the current trace. Currents were recorded at a membrane voltage of -40 mV at pH 5.5. The bar chart gives the current responses of seven individual oocytes to both substrates normalized to the sucrose response.

(b) Sucrose-induced currents plotted as a function of the corresponding sucrose concentration. Currents recorded at pH 5.5 were normalized to the maximum current measured upon application of 0.5 mM sucrose at -120 mV.

(c) Current-voltage curves recorded at pH 5.5 under 0.5 mM sucrose treatment. BvSTP13-mediated currents were normalized to the response recorded at -60 mV. Data in a-c represent means ± SEM of 7-10 individual oocytes.

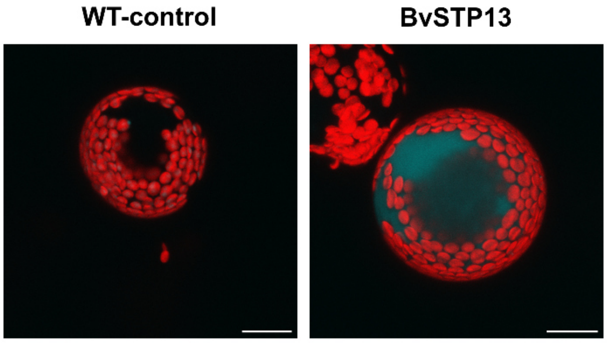


**Figure S8.** Loading of esculin in BvSTP13-transformed tobacco mesophyll protoplasts. Mesophyll protoplasts from non-transformed (left image, control) and transiently transformed *N. benthamiana leaves* (right image, BvSTP13) were incubated for 30-50 min in esculin (0.5 mM) containing buffer. After removal of residual esculin, protoplasts were analyzed with a confocal laser-scanning microscope. Z-Stacks of images from one of a total of three biological replicates were shown for control (z-stack from 5 images) and BvSTP13 (z-stacks from 7 images) transformed protoplasts. Scale bars = 20 µm.

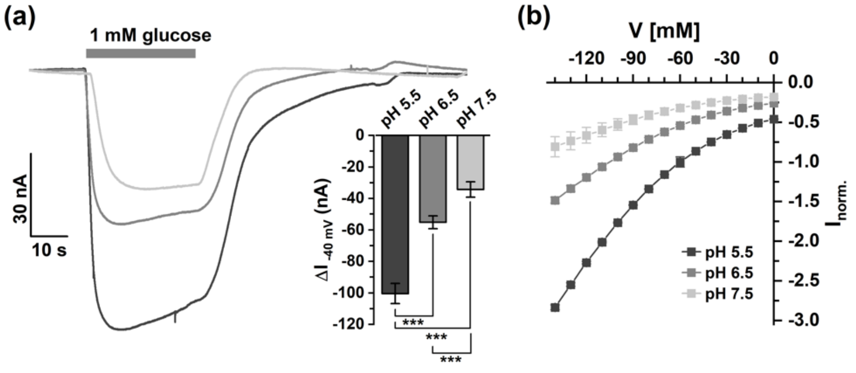


**Figure S9.** pH and voltage dependency of BvSTP13.

(a) Left panel: Representative current traces recorded from BvSTP13-expressing oocytes at the indicated pH values and a membrane voltage of -40 mV. Glucose application (1 mM) is indicated by a grey bar on the top of the current trace. Downward deflections indicate inward currents. Right panel: Glucose-induced current responses quantified for the pH conditions shown in the left panel. Data were analyzed for significant differences with paired Students *t*-test (****p* < 0.001).

(b) Current-voltage curves recorded at pH 5.5, 6.5 and 7.5 under 1 mM glucose treatment. BvSTP13-mediated current responses were normalized to the response recorded at pH 5.5 and -60 mV. Data in a (right panel) and b represent means ± SEM of 11 individual oocytes.

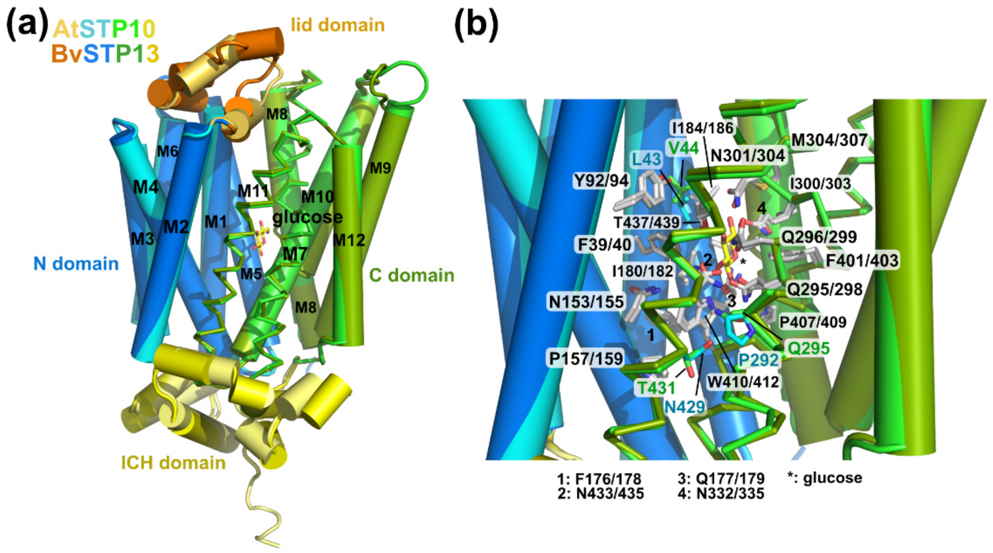


**Figure S10.** Comparison of the 3D homology model of BvSTP13 model and the crystal structure of AtSTP10.

(a) A 3D homology model of BvSTP13 was obtained based on the AtSTP10 crystal structure (wall-eye stereo view). The structures of both sugar transporters were superimposed, AtSTP10 is color-coded with light orange for the lid domain, cyan for the N domain, green for the C domain and light yellow for the C-terminal ICH domain. BvSTP13 is shown in orange for the lid domain, darker blue for the N domain, olive green for the C domain and yellow for the ICH domain. The twelve transmembrane helices are marked M1 to M12. The glucose (the bound glucose moiety of the AtSTP10 crystal structure is shown) is indicated in stick representation with the C atoms marked in yellow and the oxygen atoms in red. The lid domain differs between AtSTP10 and BvSTP13 in length and sequence, resulting in a different loop conformation. Due to an inserted proline residue, the second helix of the lid domain is split into two shorter helices in BvSTP13, which also leads to a different loop conformation for the loop between the second and third lid helix.

(b) The binding site for glucose is compared between AtSTP10 and BvSTP13 with a magnification of the binding site shown (wall-eye stereo view). Amino acid residue types identical in both AtSTP10 and BvSTP13 are marked with their carbon atoms colored in grey (nitrogen atoms in blue, oxygen atoms: red, sulfur atoms: yellow). One letter amino acid code is shown, the left number indicates the residue position in AtSTP10, the right number is the respective residue number in BvSTP13. Three amino acid residues in close proximity to the bound glucose molecule differ between AtSTP10 (residue name and number are marked in blue) and BvSTP13 (residue name and number in green). Instead of Leu43 in AtSTP10, which is placed at a distance of about 4Å to the glucose moiety, a smaller valine (Val44) occupies the same position in BvSTP13, providing slightly more space for the saccharide. At a larger distance from the glucose (about 8Å), Pro292 of AtSTP10 is replaced by glutamine (Gln295) in BvSTP13. Similarly, Asn429 of AtSTP10 is replaced by Thr431 in BvSTP13. However, the latter two residues are further away from the glucose and are located below Trp410 (Trp412 in BvSTP13), which seems to represent a bumper for the glucose moiety on its path to the cytoplasmic side, so their impact on glucose (or carbohydrate) binding is unclear.

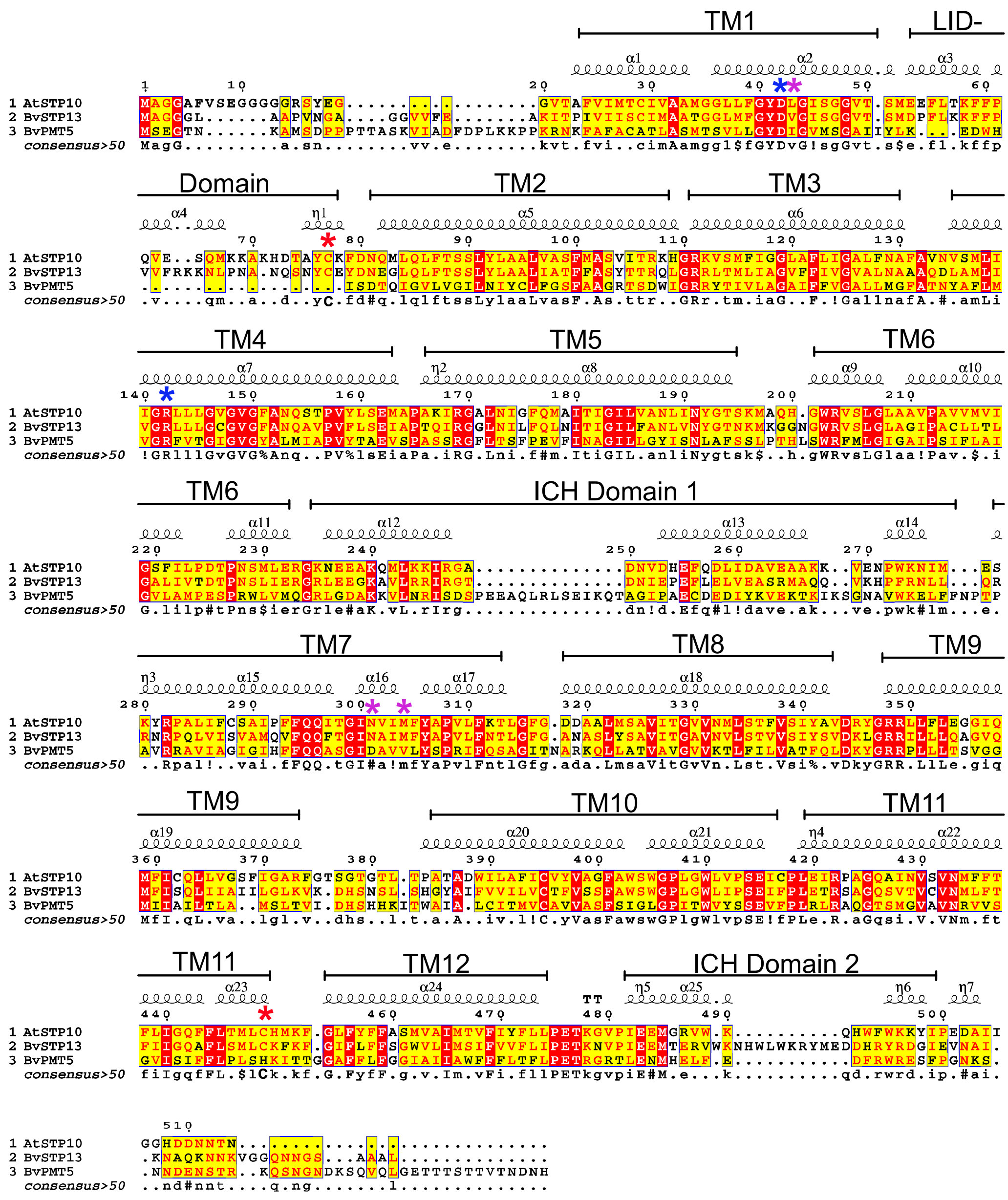

**Figure S11.** Structure guided alignment of sugar transporters. Amino acid sequences of AtSTP10, BvSTP13 and BvPMT5 were aligned using the Expresso mode of T-Coffee (Di Tommaso et al., 2011). ESPript (v3.0) (Robert & Gouet, 2014) was employed for secondary structure prediction and annotation of the alignment guided by the AtSTP10 structure (6H7D). Relative positions of the transmembrane domains (TM1 - TM12), the LID-domain and the intracellular helical bundle (ICH) domains are indicated above the sequence by barred lines. Red asterisks denote cysteine residues involved in disulfide-bridge formation between the N-terminal LID-domain and TM11 in the C-terminal of STP transporters. Blue and pink asterisks indicate the proton donor/acceptor pair needed for proton translocation and primary residues involved in sugar coordination, respectively.
